# In vivo real-time elastography with unmodified commercial endoscopes using noise-correlation-inspired method and laser speckle imaging

**DOI:** 10.64898/2026.06.23.733923

**Authors:** Maud Legrand, Nina Dufour, Francois Jonca, Jesse Schiffler, Leonardo Sosa Valencia, Nadia Bahlouli, Amir Nahas

## Abstract

Early tumor detection is critical for improving patient survival and recovery. Clinically, tissue palpation is routinely used to identify regions of abnormal stiffness, a hallmark of many pathological conditions. However, palpation is restricted to anatomically accessible sites and remains highly operator dependent. Here, we introduce a method for real-time quantitative stiffness mapping using an unmodified commercial endoscope, with the goal of enhancing diagnostic capabilities and restoring mechanical feedback during endoscopic procedures. Our approach combines shear wave elastography with speckle imaging and an innovative synchronization strategy that enables the measurement of shear wave propagation using an unmodified commercial endoscope. The resulting wave fields are analyzed with the noise-correlation-inspired (NCI) method[1], providing pixel-wise estimates of shear wave velocity and, consequently, quantitative maps of local tissue stiffness.

The method demonstrated robust performance in both benchtop and endoscopic configurations. Validation was achieved on polymer phantoms as well as on *ex vivo* and *in vivo* biological tissues, highlighting its potential for minimally invasive biomechanical imaging and real-time tissue characterization.

## 1 Introduction

Palpation is one of the oldest and most widely used methods for assessing tissue health. Because many pathological conditions, including cancer, alter tissue mechanical properties, the detection of abnormal stiffness remains a valuable diagnostic indicator[2, 3]. However, palpation is inherently qualitative, restricted to anatomically accessible regions, and strongly dependent on the practitioner’s experience. Quantitative imaging of tissue mechanics can therefore provide more objective and reproducible diagnostic information.

Elastography was introduced to address this need by mapping tissue deformation and mechanical properties[4]. Since its inception, numerous elastographic modalities have been developed, including ultrasound transient elastography[5], vibro-acoustography[6], acoustic-radiation-force-based methods[7], supersonic shear imaging[8], magnetic resonance elastography[9], and more recently optical coherence elastography[10]. These approaches have demonstrated the clinical value of stiffness imaging across a broad range of applications.

Despite these advances, translating elastography to endoscopic procedures remains challenging. Existing endoscopic elastography systems primarily rely on dedicated ultrasound probes integrated within the endoscope[11, 12]. While effective in specific clinical contexts, these systems require specialized hardware and are not easily generalized to conventional endoscopic platforms. More broadly, integrating elastography methods into commercial optical endoscopes remains a major technological challenge.

Here we introduce an elastographic imaging approach designed to operate with commercial endoscopes. Mechanical excitation is applied externally to the patient, while optical detection is performed through the endoscope camera using a laser source inserted into one of the endoscope channels. This architecture avoids major modifications of the imaging system and enables the observation of low-frequency shear waves (100–1000 Hz) within biological tissues.

Because the resulting tissue displacements are on the order of tens of nanometers, their detection requires highly sensitive optical measurements. Rather than relying on interferometric techniques such as digital holography, which are difficult to miniaturize for endoscopic applications [13], our approach exploits the sensitivity of laser speckle patterns to record small surface displacements. Speckle imaging offers a compact and robust alternative that is naturally compatible with endoscopic integration.

To recover tissue mechanical properties from the measured wave fields, we employ the Noise Correlation inspired (NCi) framework [1]. Originally derived from concepts developed in seismology and wave physics[14, 15, 16], this approach enables local shear wave velocity estimation even in configurations that do not satisfy the assumptions of conventional noise-correlation methods. By applying NCi to speckle-derived displacement measurements, quantitative stiffness maps can be reconstructed throughout the field of view. Moreover, not only the stiffness but also other properties of the tissue such as the viscosity can be accessed when combining this NCi approach with deep learning[17].

The combination of speckle-based motion detection, commercial endoscopic hardware, and NCi elastographic reconstruction provides a practical route toward real-time biomechanical imaging during minimally invasive procedures. More broadly, the framework opens the possibility of incorporating elastographic contrast into imaging systems capable of detecting small displacements, even when operating at relatively low frame rates.

## 2 Materials and methods

### 2.1 Tissue elasticity evaluation

Throughout this study, tissues were assumed to be isotropic, loss-less, quasi-incompressible, homogeneous, and semi-infinite. Their mechanical properties were characterized by the shear modulus *G*, which is related to the local shear wave velocity *c* through *G* = *ρc*^2^, where *ρ* is the tissue density. For soft biological tissues, shear wave velocities typically range from 0.5 to 10 m/s [18].

The investigated media can be approximated as semi-infinite, the waves propagating at their surface are Rayleigh waves[19]. The Rayleigh wave velocity *c*_*s*_ is related to the shear modulus by 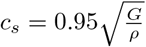, and to the wavelength *λ* and excitation frequency *f* through *c*_*s*_ = *λf*.

Assuming a tissue density close to that of water (*ρ* = 10^3^ kg*/*m^3^), local stiffness can therefore be estimated directly from the measured wave velocity.

### 2.2 From laser speckle imaging to shear wave propagation

#### Optical imaging system

Shear wave propagation was monitored using laser speckle imaging (LSI). This technique relies on the analysis of the speckle pattern generated when coherent light is backscattered by a tissue to record the propagation of shear wave.

Two LSI configurations were employed throughout this study. The first was a benchtop setup, illustrated in Fig. 1a. It consisted of a Basler a2A1920-160umBAS camera equipped with a CF12.5HA-1 Fujinon lens with an focal length of 12.5 mm. For phantom and *ex vivo* experiments, the working distance was set to 100 mm, providing a field of view of 65 *×* 36 mm, a spatial resolution of 41 *µ*m in both directions, and an acquisition rate of 200 frames per second.

**Figure 1:**
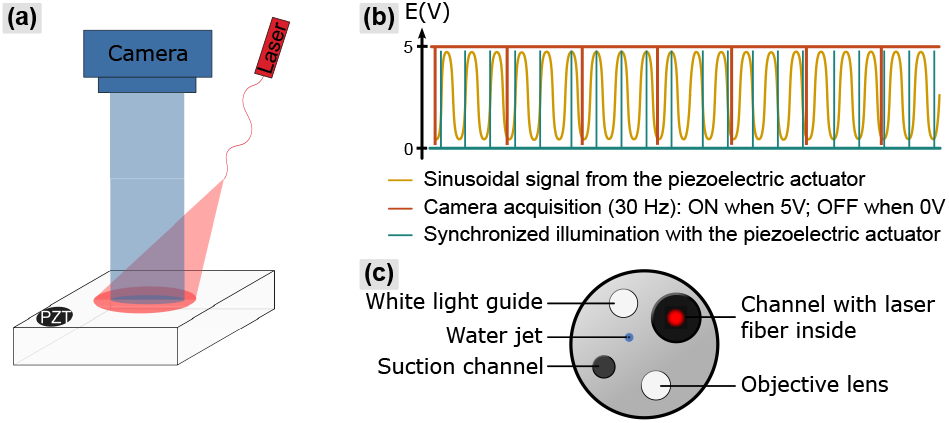
(a) Scheme of the benchtop setup and (b) the illumination synchronization. The laser diode coupled to a 300 µm multimode of 500 m long fiber emits a 637 nm beam. The piezoelectric actuator (PZT) generates a mechanical shear wave chosen between 100 and 850 Hz. To capture this propagation, the synchronization in (c) of the illumination and piezoelectric actuator signal is necessary. The sinusoidal mechanical wave (yellow) is illuminated (blue) on 4 specific values of its phase. A phase shifting of 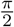 is here induced with a frequency lower than the acquisition frame rate of the camera (orange): one phase shifting is observed every two frames.

The second configuration relied on a commercial endoscope (Olympus Evis X1). A key limitation of this system was that only the video stream displayed on the monitor could be recorded, rather than the native camera output. Consequently, the available data were limited to 8-bit grayscale images acquired at 30 frames per second, which does not reflect the actual acquisition rate or performances of the endoscopic camera.

In both configurations, illumination was provided by a 637 nm laser diode (Thorlabs, L637G1). To reduce spatial coherence the beam was coupled into a 300 *µ*m-core multimode optical fiber (Thorlabs, FT300EMT) with a numerical aperture of 0.39 and a length of 500 m [20, 21]. In the benchtop configuration, the fiber was positioned as illustrated in Fig. 1a. For endoscopic measurements, the fiber was inserted into the instrument channel of the endoscope. The distal end of the endoscope with the integrated laser fiber is shown in Fig. 1c.

#### Shear wave generation

Harmonic mechanical excitation was used to generate shear waves with amplitudes on the order of a few tens of nanometers and frequencies *f* ranging from 100 to 850 Hz. Excitation was produced using a piezoelectric actuator (Cedrat Technologies, APA100M) mechanically coupled to the investigated tissue.

#### Synchronization

Synchronization is a key component of the proposed method, as it enables the observation of shear wave propagation despite the limited frame rates of the imaging systems and the unknown exposure time of the endoscopic camera.

The principle of the synchronization scheme is illustrated in Fig. 1b. The laser illumination is synchronized with the piezoelectric actuator such that only a small fraction of the mechanical cycle is illuminated, corresponding to less than one twentieth of the wave period. As a result, each illumination pulse samples a specific phase of the propagating shear wave.

Because the mechanical excitation frequency is substantially higher than the camera acquisition frequency, only a limited number of photons would be collected if a single illumination pulse were used during each camera exposure. To increase the detected optical signal, the same phase of the mechanical wave is illuminated repeatedly during a single camera exposure time. This strategy accumulates photons corresponding to an identical mechanical state and thereby increases the signal associated with a given wave phase during one exposure time.

To reconstruct shear wave propagation, several distinct phases of the wave must be sampled. Although the Noise Correlation inspired (NCi) method requires a minimum of three phases, four phases were used throughout this study to improve robustness against noise. Consequently, only four discrete phases of the mechanical wave are acquired rather than the full wave propagation sequence.

The transition between successive phases is achieved by introducing a temporal delay in the illumination sequence. Every two camera frames, the illumination timing is shifted by *π/*2 relative to the previous phase. Although the camera itself is not synchronized to the illumination sequence, its acquisition frequency is known, allowing the phase shift to be applied deterministically throughout the acquisition.

As a consequence, every two recorded frames contain information from a single wave phase, while the intermediate frames contain a mixture of two successive phases due to the temporal shift. The exploitation of these single-phase and mixed-phase frames is described in the following section.

Importantly, this synchronization strategy provides an alternative to the use of high-speed cameras that would otherwise be required to directly resolve shear wave propagation. It therefore enables elastographic imaging using low frame rate or low-cost or uncontrolled imaging systems.

#### Speckle image processing

Shear wave propagation was extracted from the recorded speckle movie using a processing approach inspired by the work of Fujii *et al*. [22]. For two consecutive images *I*_*k*_ and *I*_*k*+1_, the speckle contrast variation was calculated as:

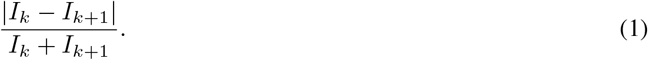

As described above, only one frame out of two corresponds to a unique phase of the mechanical wave, whereas the intermediate frames contain a mixture of two phases. Consequently, the speckle processing was performed using only the frames associated with unique phases.

The resulting signal was spatially filtered using a Gaussian blur with a standard deviation of 5 pixels before further analysis.

### 2.3 From shear wave propagation to real-time elastography

The conversion of the measured shear-wave propagation into quantitative stiffness maps relies on the Noise Correlation inspired (NCi) framework [1, 23, 24]. This approach establishes a relationship between the measured displacement field, the impulse response of the medium (ie. the Green’s function), and the spatial-temporal correlation function, enabling the estimation of local shear-wave velocity.

When a mechanical source *s*(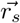, *t*) generates a shear wave at position 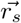, the axial displacement *ψ*_*z*_ measured at position 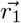 can be expressed as: 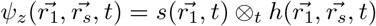. where ⊗_*t*_ denotes temporal convolution and *h*(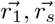, *t*) is the impulse response of the medium from 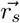 to 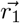.

Considering two positions 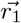 and 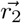, the definition of the spatial correlation is: 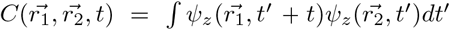. This quantity is equal to the correlation with a time-reversed function: 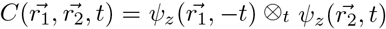. Using the reciprocity principle[24], the correlation of the impulse responses can be written: 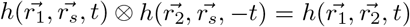. By defining *S*(*t*) = *s*(*t*) ⊗_*t*_ *s*(*−t*) which corresponds to the source autocorrelation, the spatial correlation becomes:

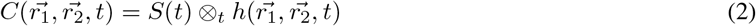

Under the assumptions described above (homogeneous, isotropic, lossless, semi-infinite medium), the impulse response is a Dirac distribution. Thus, the equation 7 evaluated in the spatial and temporal auto-correlation gives:

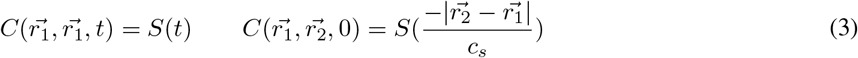

These relations show that the local shear-wave velocity *c*_*s*_ can be estimated from the spatial and temporal correlation functions if the shear-wave source is known.

#### Full width at half maximum method

In order to estimate the shear-wave velocity, a first approach consists of estimating the spatial and temporal full width at half maximum (FWHM) of the correlation functions defined in Eq. 3.

The ratio between these two quantities provides an estimate of the shear-wave velocity:

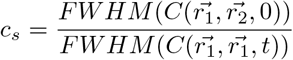

Although straightforward to implement, this method requires the calculation of complete spatial and temporal correlation curves at every image pixel, resulting in increased computational cost and low spatial resolution (about the size of the shear-wave wavelength).

#### Noise Correlation inspired (NCi) method

To overcome these limitations, the Noise Correlation inspired (NCi) method has been developed [1]. Rather than relying on the full correlation function, NCi only requires few points at the top of it to extract the second order derivative, which is calculated using the Hessian matrix, defined as:

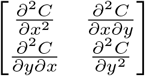

From the first eigenvalue of this hessian matrix, the velocity can be retrieved. Indeed, as demonstrated by Marmin *et al*.[25], the second-order temporal and spatial derivatives of the correlation function can be directly related to the local shear-wave velocity through

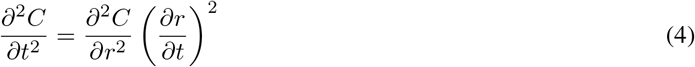

Thus,

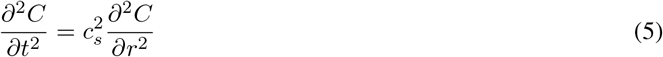

This relationship can be rewritten as

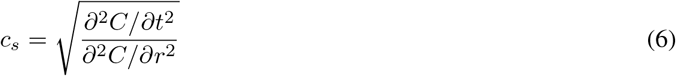

showing that the local shear-wave velocity can be estimated directly from the ratio of the second-order temporal and spatial derivatives of the correlation function.

In practice, the spatial curvature is obtained from the first eigenvalue of the Hessian matrix evaluated around the autocorrelation peak. Consequently, only a small neighborhood surrounding the autocorrelation maximum is required to estimate the velocity. Typically, 5 to 10 neighboring pixels are sufficient, enabling pixel-wise velocity estimation while preserving a high spatial resolution.

Throughout this study, the FWHM method was used as an independent reference to validate the velocity estimates obtained with the NCi method.

#### Real-time implementation

The mechanical shear wave that propagates is harmonic and can therefore be written in each pixel as *cos*(*ωt − ϕ*) with *ϕ* an arbitrary initial phase. Regardless of the value of *ϕ*, the autocorrelation of a harmonic signal remains proportional to cos(*ωt*) and exhibits a maximum at *t* = 0.

Since the temporal autocorrelation is known to be proportional to cos(*ωt*), the Nyquist-Shannon sampling theorem implies that only three samples per period are theoretically required to reconstruct it. In practice, four equally spaced phases were acquired to improve robustness against noise: *cos*(*ωt − ϕ*), 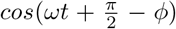, *cos*(*ωt* + *π − ϕ*) and 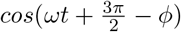. These four synchronized phases are illustrated in Fig. 1b.

Consequently, the temporal autocorrelation can be reconstructed from only four phase images, enabling the estimation of local wave velocity and tissue stiffness without acquiring the complete wave propagation sequence. Combined with the synchronization strategy described above, this approach enables real-time stiffness mapping, even when using the 30-Hz video stream provided by the commercial Olympus Evis X1 endoscope.

### 2.4 Simulation of speckle-modulated shear-wave propagation

Finite-element simulations were performed to validate the combination of the speckle-processing pipeline and the Noise Correlation inspired (NCi) method described above. The simulations were designed to reproduce the experimental conditions as closely as possible, with particular emphasis on the gelatin phantom experiments.

Gelatin samples were modeled as homogeneous parallelepipeds with dimensions of 65 *×* 65 *×* 20 mm. The material was assumed to be isotropic, linear elastic, and nearly incompressible. Nearly incompressible behavior was modeled by setting the Poisson ratio to *v* = 0.495. To avoid shear-locking effects, the volume was meshed using Gmsh with 10-node tetrahedral elements, resulting in a mesh composed of 225,927 nodes and 162,560 elements.

Boundary conditions were selected to reproduce the experimental setup. Zero displacement along the *z* direction was imposed on the bottom surface (*h* = 0 mm), corresponding to the contact with the supporting plate. To suppress rigid-body motion, zero displacement along the *x* and *y* directions was additionally imposed on lines of the bottom surface. Mechanical excitation was applied on the upper surface (*h* = 10 mm) over a circular area matching the diameter and position of the piezoelectric pin used experimentally. The imposed displacement was defined as:

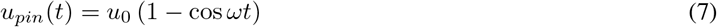

The simulations were performed using FenicsX [26, 27, 28, 29] on a MacBook Air equipped with an Apple M4 processor (10 cores, 16 GB RAM). Seven CPU cores were used through OpenMPI parallelization. The simulated duration was 48 ms with 200 temporal iterations. All simulations converged successfully, with computation times below 1 h 44 min.

Several shear moduli were investigated (*G* = *{*4, 8, 16, 25, 36, 49, 64*}* kPa). Only the out-of-plane displacement omponent was retained for subsequent analysis and interpolated onto a regular 500 *×* 500 grid using bicubic interpolation.

The simulated displacement fields were subsequently combined with synthetic speckle patterns using MATLAB R2023b. Speckle was modeled through random phase modulation of the optical field. Because the displacement amplitudes under investigation are on the order of tens of nanometers, they were assumed to induce only a phase modulation of the speckle field while leaving its amplitude unchanged. The resulting signal *S* was expressed as

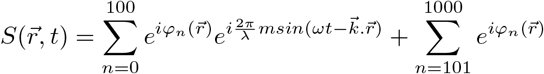

where:

1. *m* denotes the displacement amplitude (100 nm).
2. φ_*n*_ represents the random phase added.
3. *λ* denotes the laser wavelength (600 nm).
4. *ω* = 2*πf*, where *f* is the excitation frequency.

The first term represents scatterers affected by the propagating shear wave, while the second term corresponds to static speckle contributions. An additive white uncorrelated noise was subsequently introduced, corresponding to a signal-to-noise ratio (SNR) of 10.

The temporal evolution of φ_*n*_ was neglected in these simulations. This approximation assumes that the tissue remains static or quasi-static over the time scale of shear wave propagation.

The simulated speckle sequences were processed using the same speckle-processing pipeline and NCi method as the experimental data. The resulting velocity maps are presented in the Results section.

### 2.5 Sample preparation

Three types of samples were investigated throughout this study: gelatin phantoms, *ex vivo* pork liver tissue, and *in vivo* human tissues.

Gelatin phantoms measured 65 *×* 65 *×* 20 mm. They consisted of 1 wt% titanium dioxide, 10 to 30 wt% pork gelatin, and distilled water. *Ex vivo* liver samples were obtained from pork liver sections approximately 10 *×* 5 *×* 2 cm in size. Samples were either used in their native state or modified to include a localized stiff inclusion. The inclusion was created by partial thermal coagulation using a high-intensity pulsed laser (Amplitude Satsuma X), resulting in a roughly cylindrical region approximately 1.5 cm in diameter and 1 cm in depth.

For the *in vivo* experiments, measurements were performed on the forearm and wrist of healthy volunteers. All procedures were conducted in accordance with the Declaration of Helsinki, and written informed consent was obtained from all participants prior to the experiments.

## 3 Results

The proposed framework was evaluated through numerical simulations, benchtop experiments, and endoscopic measurements. Validation was performed progressively on simulated wave fields, tissue-mimicking phantoms, *ex vivo* and *in vivo* biological tissues, and finally using a commercial endoscope. Two experimental systems were used: a benchtop setup, and a commercial endoscopic setup. Additional details regarding the experimental configurations are provided in the **Materials and Methods** section.

### Simulated data

To validate the method under controlled conditions, monochromatic shear wave propagation was simulated over a 50 mm field of view with a spatial sampling of 100 *µ*m and a mechanical excitation frequency of 500 Hz. To reproduce experimental conditions, the simulations included realistic simulated speckle patterns as well as additive white noise corresponding to an SNR of 10, matching the experimental conditions. Seven stiffness values ranging from 4 to 64 kPa were simulated, and each simulation was repeated five times.

The speckle-processing pipeline described in the section **Materials and Methods** successfully recovered the underlying shear wave propagation from the simulated speckle movies (Fig.2a). The resulting wave fields were analyzed using the NCi method to estimate local wave velocity (Fig.2b). For a simulated velocity of 7 m/s, the reconstructed median velocity was also approximately 7 m/s, with a standard deviation of about 2.5 m/s. Across all seven stiffness values, reconstructed shear moduli closely matched the theoretical values (Fig.2c). Linear regression between expected and reconstructed stiffness yielded a correlation coefficient of 0.9991, demonstrating the ability of the proposed framework to accurately recover tissue mechanical properties from simulated wave fields movies.

**Figure 2:**
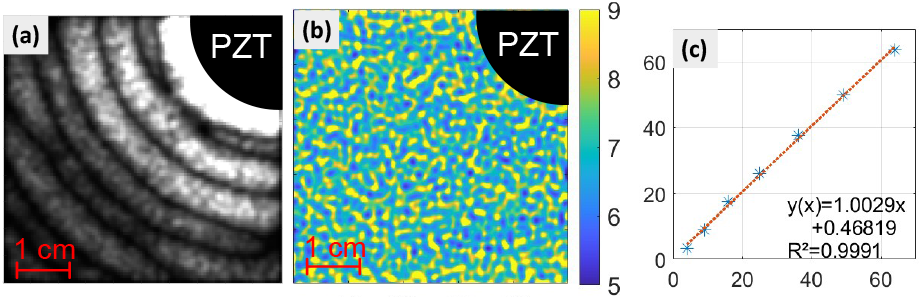
Simulated shear wave propagation for different shear modulus. (a) Shear wave propagation image with speckle pattern, after applying the speckle image processing for a 500 Hz wave and a stiffness of 49 kPa. (b) Associated velocity map (*m*.*s*^*−*1^) retrieved with Noise Correlation inspired method. (d) Comparison of the median shear modulus values found with the simulation, and the theoretical values. PZT: piezoelectric actuator

### Quantitative stiffness mapping in gelatin phantoms

Following validation on simulated data, the method was evaluated experimentally using gelatin phantoms (Louis Francois Ingrédients Alimentaires 735H, Pork Gelatin powder 200°Bloom) with controlled optical and mechanical properties. Titanium dioxide particles (Sigma-Aldrich, CAS no.1317-70-0, -325 mesh powder) were added to generate optical scattering conditions comparable to those encountered in biological tissues[1]. Mechanical properties were adjusted by varying the gelatin concentration from 10% to 30%. Representative results obtained on a 10% gelatin phantom are shown in Fig.3. Shear wave propagation generated by a 700 Hz excitation was clearly recovered after speckle processing (Fig. 3a). Application of the NCi method yielded a velocity map with a median velocity of approximately 1.8 m/s in the high-SNR region *α* (Fig.3b). Although illumination artifacts are visible in the processed wave-field images (red arrows in Fig.3a), these regions do not affect velocity estimation. The local speckle dynamics induced by wave propagation remain measurable, allowing accurate velocity reconstruction despite non-homogeneous image intensity. Consequently, these artifacts are not visible in the final velocity map. In contrast, regions characterized by low signal-to-noise ratios (SNR *<* 2, region *β* in Fig.3b) exhibited larger velocity errors, higher than 0.8 m/s relative to the values measured in the high-SNR region. To independently assess accuracy, the wavelength was manually measured directly from the propagation images. This analysis yielded a velocity of approximately 1.6 m/s, in close agreement with the 1.8 m/s median value estimated by the NCi method. The resulting absolute error remained below 10%, which is comparable to values commonly reported in shear wave elastography studies[30, 1]. The influence of phantom stiffness was further investigated using gelatin concentrations ranging from 10% to 30% and excitation frequencies between 600 and 850 Hz (Fig.3c). For each concentration, at least ten independent measurements were performed. Increasing gelatin concentration systematically increased wave velocity, consistent with the expected increase in stiffness, demonstrating the robustness and repeatability of the measurements. Furthermore, only limited velocity variations were observed across the investigated frequency range, demonstrating the expected low viscoelasticity of gelatin.

**Figure 3:**
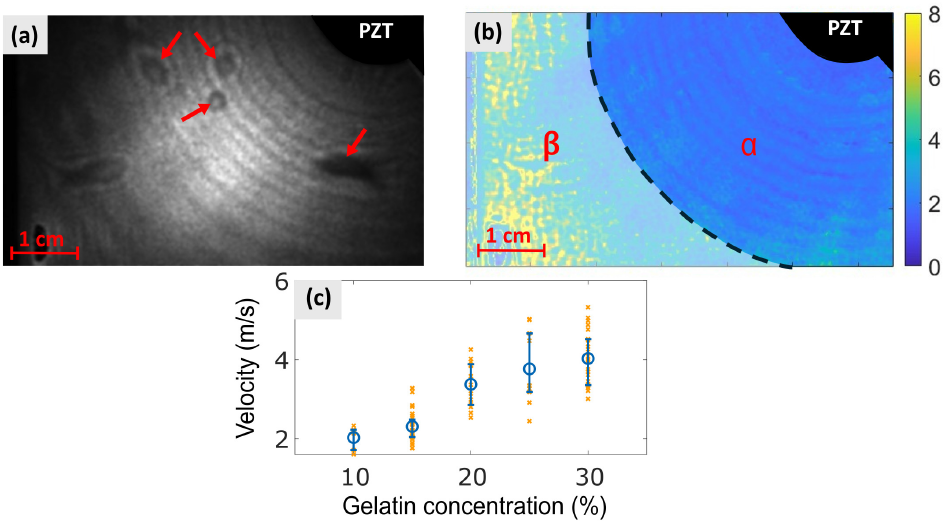
Shear modulus for different frequencies and 4 different gelatin concentration samples. (a) Shear wave propagation image after speckle image processing for a 700 Hz sinusoidal wave on a 10% gelatin phantom. (b) Velocity map (*m*.*s*^*−*1^) retrieved with NCi method. (c) Velocity values (*m*.*s*^*−*1^) for different gelatin concentration, with mechanical excitation frequency between 600 and 850 Hz. PZT: piezoelectric actuator

### *Ex vivo* and *in vivo* biological tissues

To evaluate the applicability of the method to biological tissues and conditions closer to future endoscopic applications, experiments were performed on *ex vivo* pork liver and *in vivo* human tissues using the benchtop setup. These experiments therefore provide a rigorous test of the robustness of the proposed approach under realistic imaging conditions.

Two configurations were investigated for the *ex vivo* pork liver. The first consisted of a homogeneous raw liver sample (Fig.4a-b). The second was designed to mimic a stiff inclusion embedded within a softer surrounding tissue, analogous to a tumor within healthy tissue. This heterogeneous configuration was obtained by introducing a cooked inclusion into the raw liver (lighter region in Fig.4c-d). Additional details regarding sample preparation are provided in the **Materials and Methods** section.

For the homogeneous liver sample, shear-wave propagation generated by a 100 Hz excitation was successfully recovered after speckle processing (Fig.4a). The corresponding velocity map is shown in Fig.4b. Reliable measurements were obtained in the vicinity of the piezoelectric actuator, where the wave amplitude remained sufficiently high. In this region, the median velocity was approximately 1 m/s, corresponding to a shear modulus of about 1 kPa according to the assumptions detailed in the **Materials and Methods** section. This value is consistent with previously reported measurements obtained at similar excitation frequencies, which yielded shear moduli slightly above 2 kPa [31].

The heterogeneous liver sample further demonstrated the ability of the method to detect local stiffness variations (Fig.4c-d). The cooked inclusion was clearly visible in the reconstructed velocity map as a region of elevated wave velocity. The inclusion exhibited a median velocity of approximately 1.8 m/s, corresponding to a shear modulus of about 3.5 kPa, whereas the surrounding raw tissue displayed a median velocity close to 1 m/s and a shear modulus of approximately 1 kPa. To the best of our knowledge, no reference values are available for cooked pork liver under these experimental conditions. Nevertheless, the measured stiffness increase is consistent with the expected mechanical contrast between cooked and uncooked tissue and remains compatible with previously reported values for raw liver [31]. Regions located further from the actuator exhibited lower reliability due to several combined effects, including reduced illumination caused by tissue uneven surface and attenuation of the propagating shear wave. In these regions, the signal-to-noise ratio dropped below 2, limiting the accuracy of the reconstructed velocities.

The method was subsequently evaluated *in vivo* on the forearm and wrist, at 250 Hz. The forearm provides a relatively homogeneous tissue environment, whereas the wrist contains a more complex arrangement of stretched skin, muscle, and tendon structures. For the forearm measurements (Fig. 4e-f), the median shear modulus was approximately 1.4 kPa, corresponding to a wave velocity of about 1.2 m/s. An independent manual wavelength measurement yielded a velocity close to 1 m/s, in good agreement with the value estimated using the NCi framework.

**Figure 4:**
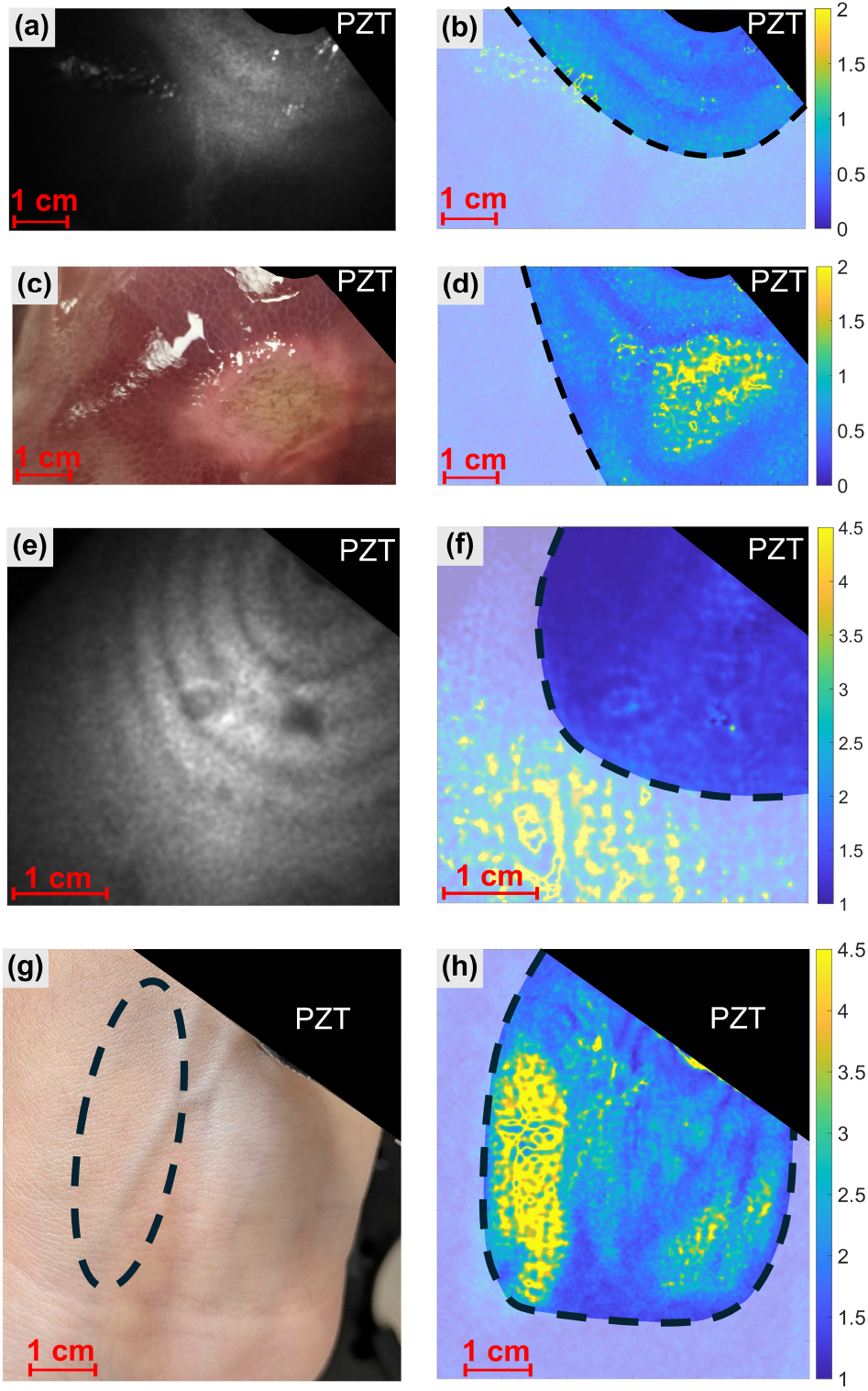
Acquisitions and velocity map retrieved on samples. (a) is the raw image of an *ex vivo* pork liver with a round cooked inclusion inside, excited at 100 Hz. (b) is the associated calculated velocity map (*m*.*s*^*−*1^). (c) is the speckle-processed image of an *ex vivo* homogeneous pork liver to highlight the shear wave propagation inside an homogeneous stiffness area, at 100 Hz. (d) is the associated calculated velocity map (*m*.*s*^*−*1^). (e) is the speckle-processed image of an *in vivo* forearm excited at 250Hz with (f) its associated velocity map. (g) is the raw image of the inside part of a wrist, with a tendon (surrounded) under a stretched skin at 500 Hz, with (h) its velocity map. PZT: piezoelectric actuator

More pronounced mechanical heterogeneity was observed in the wrist measurements (Fig. 4g-h). The region corresponding to stretched skin and muscle exhibited a shear modulus of approximately 16 kPa, associated with a wave velocity of about 4 m/s. Independent manual wavelength measurement yielded a velocity close to 3 m/s. The tendon region appeared significantly stiffer, with a shear modulus of approximately 30 kPa and a corresponding velocity of about 5.5 m/s. Manual wavelength measurement provided a velocity estimate of approximately 4.5 m/s. Overall, the measured mechanical contrasts were consistent with the known relative stiffness of the investigated anatomical structures and showed good agreement with independently estimated wave velocities.

### Commercial endoscope configuration

The ultimate objective of this work is to enable quantitative elastography using commercially available endoscopes. To assess the feasibility of this approach, the complete framework was implemented in a commercial endoscopic configuration and evaluated on both gelatin phantoms and *in vivo* human tissue (Fig. 5).

**Figure 5:**
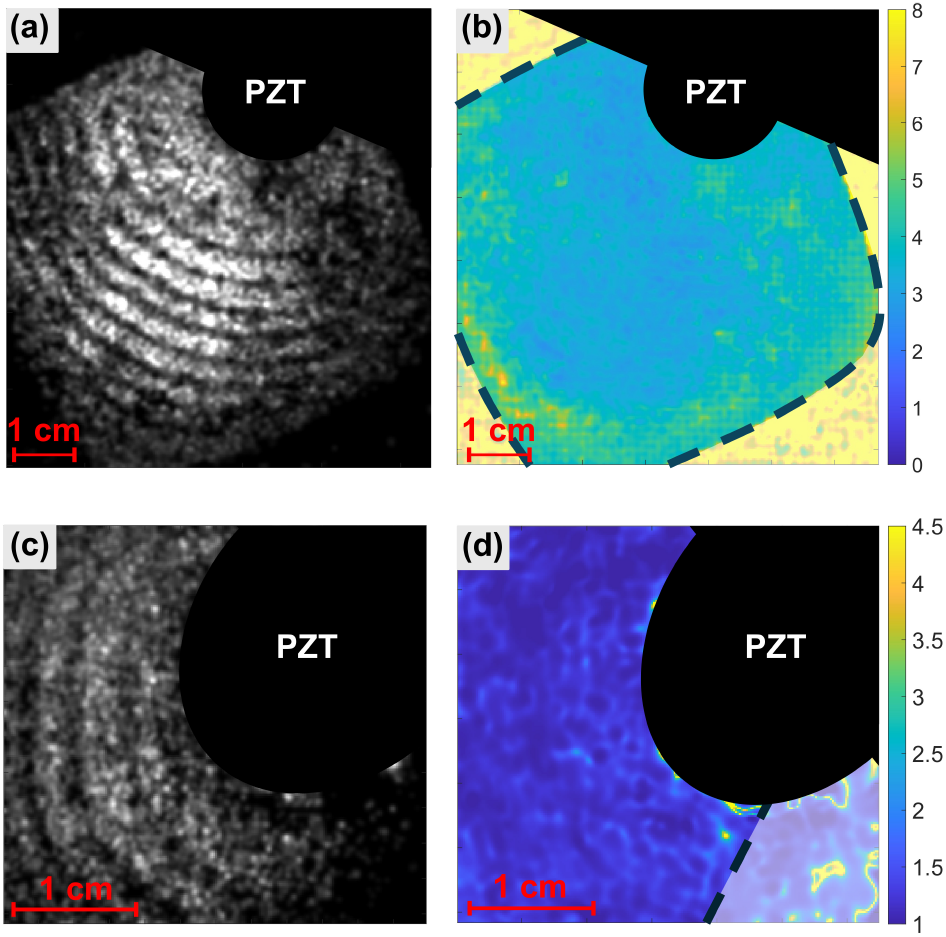
Acquisitions made with the endoscopic setup and associated velocity maps. (a) is a 650 Hz shear wave propagating inside a 20% gelatin sample, and (b) the velocity map (*m*.*s*^*−*1^) retrieved from the NCi method, observed with the endoscopic camera. (c) corresponds to an *in vivo* forearm with a 250 Hz shear propagation, with (d) its associated velocity map (*m*.*s*^*−*1^). PZT: piezoelectric actuator

A first validation was performed on a 20% gelatin phantom using a 650 Hz mechanical excitation (Fig. 5a-b). Shearwave propagation was successfully recovered from the endoscopic images, and application of the NCi method yielded a median wave velocity of approximately 3.8 m/s. This corresponds to a shear modulus of about 15 kPa. Importantly, this value is in close agreement with the measurements obtained using the benchtop setup for gelatin samples of identical concentration (Fig. 3c), demonstrating that the transition from a laboratory imaging system to a commercial endoscope does not significantly affect stiffness estimation.

The performance of the system was subsequently evaluated *in vivo* on the forearm using a 250 Hz mechanical excitation (Fig. 5c-d). The reconstructed velocity map yielded a median velocity of approximately 1.5 m/s. This value is consistent with the velocity previously measured on the forearm using the benchtop configuration (Fig. 4f), further confirming the reliability of the proposed approach under endoscopic imaging conditions.

Together, these results demonstrate that quantitative stiffness mapping can be achieved using a commercial endoscope without major modifications of the imaging system. The close agreement between benchtop and endoscopic measurements highlights the robustness of the proposed framework and supports its potential for real-time biomechanical characterization during minimally invasive procedures.

Overall, the proposed framework consistently recovered tissue mechanical properties across simulations, tissue-mimicking phantoms, and biological tissues. Quantitative stiffness estimations remained consistent with independent measurements and literature values, while preserving sensitivity to local mechanical heterogeneities. Most importantly, comparable results were obtained using both benchtop and commercial endoscopic configurations, demonstrating the feasibility of real-time elastographic imaging with minimally modified clinical instrumentation.

## 4 Discussion

In this study, we demonstrate that the Noise Correlation inspired (NCi) method can be combined with speckle imaging to perform quantitative elastography using a commercial endoscope. By coupling optical detection of shear wave propagation with an innovative synchronization strategy, the proposed approach enables local stiffness estimation using standard endoscopic imaging hardware.

The methodology was first validated using numerical simulations. The calculated velocity and stiffness values showed excellent agreement with the expected mechanical properties, confirming the ability of the combined speckle-processing and NCi method to recover local wave propagation. Although the simulated stiffness values were accurately reconstructed, further improvements could be achieved by refining both the noise model and the representation of speckle in the simulations. In particular, in the present implementation, shear wave propagation is introduced as a phase modulation of the speckle pattern, consistent with the multiplicative nature of speckle noise. More advanced models may provide a more realistic representation of the experimental conditions and further improve the accuracy of numerical predictions.

Experimental validation was subsequently performed on gelatin phantoms with controlled optical and mechanical properties. Quantitative velocity measurements were obtained with errors below 10% when compared with independent wavelength-based estimates, demonstrating the accuracy of the proposed approach. Repeated measurements further confirmed the reproducibility of the method. The observed variability between acquisitions is likely attributable to temperature-dependent changes in gelatin stiffness, a phenomenon that has previously been reported in the literature[32].

The method was then evaluated on *ex vivo* and *in vivo* biological tissues. Quantitative velocity estimates were successfully recovered in all investigated tissues. In addition, the method was able to detect local mechanical heterogeneities, including the stiff inclusion embedded within the pork liver sample and the distinct stiffness contrast between soft tissue and tendon structures in the wrist. These results demonstrate that the method remains effective under realistic biological conditions despite increased tissue complexity.

The most significant result of this study is the successful implementation of the whole method *in vivo* using a commercial endoscope. Measurements performed on both gelatin phantoms and *in vivo* tissues yielded velocity values consistent with those obtained using the benchtop setup, demonstrating that quantitative elastography can be achieved without specialized endoscopic imaging hardware. This compatibility with commercially available devices represents an important step toward the integration of biomechanical imaging into routine minimally invasive procedures.

Notably, all endoscopic measurements were obtained from screen recordings rather than directly from the camera output. Consequently, the reconstructed wave fields were generated from data limited to 8-bit grayscale levels and a frame rate of only 30 Hz. Despite these constraints, reliable stiffness estimates were recovered. These results suggest that the proposed framework remains robust even under imaging conditions that are substantially less favorable than those typically required for conventional shear-wave imaging.

A key component of the proposed framework is the synchronization strategy described in the **Materials and Methods** section. Thanks to this approach, only four fixed phases of the shear wave need to be illuminated and recorded. Although the camera is unable to directly resolve wave propagation, the same wave phase is illuminated repeatedly and accumulated within each frame, providing sufficient optical signal for wave reconstruction. Consequently, quantitative elastography can be performed using low-cost or uncontrolled imaging systems that would normally be unable to track shear wave propagation. More broadly, this strategy opens the possibility of incorporating elastographic contrast into any imaging systems capable of detecting small displacements, even when operating at low frame rates.

Finally, because only four images are required to reconstruct the elasticity map, not only the measurement is *in vivo* compatible but also the computational time remains low and is compatible with real-time implementation. Together, these results establish a practical framework for quantitative endoscopic elastography and provide a pathway toward real-time biomechanical characterization using commercially available imaging platforms.

## Data Availability Statement

The data that support the findings of this study are available from the corresponding author upon reasonable request.

## Acknowledgments

This work would have not been the same without the help of few people. Q. Becar (ICube Laboratory) for his insightful discussions on laser diodes, B. De Azevedo (ICube Laboratory) for the cooking of pork liver and A. Gressier (IHU Strasbourg) who gently gave us a large access to the endoscope. This work was supported by Agence Nationale de la Recherche (No. ANR-21-CE19-0018-01), Interdisciplinary Thematic Institute HealthTech (No. ANR-10-IDEX-0002 and STRAT’US Project No. ANR-20-SFRI-0012), France Life Imaging (No. ANR-11-INBS-0006), and the Region Grand Est.

